# ATP-Binding Cassette Family C member 1 constrains metabolic responses to high-fat diet in male mice

**DOI:** 10.1101/2024.01.23.576896

**Authors:** Elisa Villalobos, Allende Miguelez-Crespo, Ruth A. Morgan, Lisa Ivatt, Dominic Kurian, Judit Aguilar, Rachel A. Kline, Thomas M. Wishart, Nicholas Morton, Roland H. Stimson, Ruth Andrew, Brian R. Walker, Mark Nixon

**Affiliations:** University/British Heart Foundation Centre for Cardiovascular Science, The Queen’s Medical Research Institute, University of Edinburgh, Edinburgh, United Kingdom, EH16 4TJ; Scotland’s Rural College, The Roslin Institute, Easter Bush Campus, United Kingdom; The Roslin Institute, Royal (Dick) School of Veterinary Studies, College of Medicine and Veterinary Medicine, University of Edinburgh, Easter Bush, United Kingdom; Translational and Clinical Research Institute, Newcastle University, Newcastle upon Tyne, United Kingdom; Centre for Systems Health and Integrated Metabolic Research, Nottingham Trent University, Nottingham, United Kingdom

**Keywords:** Glucocorticoids, steroids, transport, metabolism, obesity, homeostasis

## Abstract

Glucocorticoids modulate glucose homeostasis, acting on metabolically active tissues such as liver, skeletal muscle, and adipose tissue. Intra-cellular regulation of glucocorticoid action in adipose tissue impacts metabolic responses to obesity. ATP-Binding Cassette Family C member 1 (ABCC1) is a transmembrane glucocorticoid transporter known to limit the accumulation of exogenously administered corticosterone in adipose tissue. However, the role of ABCC1 in the regulation of endogenous glucocorticoid action and its impact on fuel metabolism has not been studied. Here, we investigate the impact of *Abcc1* deficiency on glucocorticoid action and high fat-diet (HFD)-induced obesity. In lean mice, deficiency of *Abcc1* increased endogenous corticosterone levels in skeletal muscle and adipose tissue but did not impact insulin sensitivity. In contrast, *Abcc1*-deficient mice on HFD displayed impaired glucose and insulin tolerance, and fasting hyperinsulinemia, without alterations in tissue corticosterone levels. Proteomics and bulk RNA sequencing in adipose tissue and skeletal muscle revealed that *Abcc1* deficiency amplified the transcriptional response to an obesogenic diet in adipose tissue. Moreover, the *Abcc1* deficiency impairs key signalling pathways related to glucose metabolism in both skeletal muscle and adipose tissue, in particular those related to OXPHOS machinery and Glut4. Together, our results highlight a role for ABCC1 in regulating glucose homeostasis, demonstrating diet-dependent effects that are not associated with altered tissue glucocorticoid concentrations.

## INTRODUCTION

Glucocorticoids are required to maintain glucose and lipid homeostasis in times of physiological stress, ensuring an adequate fuel supply for the body (Kuo, et al. 2015). In key metabolic tissues such as adipose tissue, skeletal muscle and liver, glucocorticoids act to prevent glycolysis and instead promote gluconeogenesis (Kuo, et al. 2013). However, chronic glucocorticoid excess, e.g. in Cushing’s syndrome, causes metabolic dysfunction including hyperglycaemia and obesity (Kaikaew, et al. 2019; Pivonello, et al. 2016). Moreover, increased tissue levels of glucocorticoids have been described in obese humans and in animal models of obesity (Anderson, et al. 2016; Luijten, et al. 2019; Pivonello et al. 2016). In particular, increased adipose (Gathercole, et al. 2007; Kershaw, et al. 2005; Rask, et al. 2001) or skeletal muscle (Loerz and Maser 2017; Morgan, et al. 2009; Trost, et al. 2002) exposure to glucocorticoids is accompanied by insulin resistance, consistent with the effects of glucocorticoids on glucose and lipid utilisation.

Controlling intracellular levels of glucocorticoids within target tissues can influence plasma glucocorticoid levels over and above their regulation by the hypothalamic-pituitary-adrenal (HPA) axis. Studies have shown that removing the effect of glucocorticoids, e.g. by adrenalectomy or inhibition of 11-beta-hydroxysteroid dehydrogenase type 1 (11β-HSD1), prevents weight gain and metabolic dysfunction in animal models of obesity and in humans (Anderson et al. 2016; Bray, et al. 1992; Gambineri, et al. 2014; Sainsbury, et al. 1997). However, 11β-HSD1 inhibitors have not progressed beyond phase II trials due to their insufficient efficacy on metabolic outcomes (Rosenstock, et al. 2010; Scott, et al. 2014). We have also recently shown that mice with deficiency of carbonyl reductase 1 (*Cbr1*), a further glucocorticoid regulator in adipose tissue, display lower levels of fasting glucose and improved glucose tolerance (Bell, et al. 2021). A key question is whether there are other mechanisms that confer tissue-specific control of glucocorticoid levels that might be important in obesity and tractable to therapy.

ATP-Binding Cassette Subfamily C Member 1 (ABCC1) is a multidrug efflux transporter present in the plasma membrane of many cell types, including adipocytes and myotubes, but not hepatocytes (Ling, et al. 2014; Meijer, et al. 1998; Qu, et al. 2014; Sajja and Cucullo 2015; Shi, et al. 2017). *ABCC1* is highly expressed in adipose tissue and skeletal muscle, but poorly expressed in liver (Devine, et al. 2023). While primarily studied due to its capacity to efflux drugs and its role in determining resistance to chemotherapeutic agents in cancer (Bernal-Sore, et al. 2018; Ling et al. 2014; Meijer et al. 1998; Mohankumar, et al. 2018; Shi et al. 2017), we recently identified ABCC1 as a regulator of HPA-axis negative feedback in humans (Kyle, et al. 2022), and as a glucocorticoid transporter in white adipose tissue (Nixon, et al. 2016). Mice with global deletion of *Abcc1* or mice administered the ABCC1 inhibitor, probenecid, accumulate exogenously-administered corticosterone within adipose tissue, associated with exaggerated subcutaneous adipose glucocorticoid-responsive gene transcription. Moreover, analysis of *ABCC1* expression revealed an increase in mRNA levels in subcutaneous and visceral adipose tissue from obese patients compared with lean controls (Nixon et al. 2016), suggesting a compensatory mechanism to ‘protect’ adipose from glucocorticoid excess. Additionally, recent evidence demonstrates relatively high levels of *ABCC1* in skeletal muscle in humans and mice (Devine et al. 2023; Nixon et al. 2016), indicating a potential role for this efflux transporter in regulating glucocorticoid action in key extra-adipose metabolic tissues. Importantly, the role of *Abcc1* in murine models of obesity has not yet been described.

We tested the hypothesis that mice lacking *Abcc1* exhibit an adverse metabolic profile due to increased intra-tissue endogenous glucocorticoid action in white adipose tissue and skeletal muscle. Using a global knockout of *Abcc1* in adult male mice we aimed to evaluate the influence of *Abcc1* on the metabolic profile in both lean and obese conditions, using control chow and high fat diet, respectively.

## MATERIALS AND METHODS

### Animals

Male *Abcc1* knockout (*Abcc1*-KO) mice were purchased from The Jackson Laboratory (B6.129S1-Abcc1tm1Acs/VoreJ, Stock No: 028129) and bred in house with female C57BL/6J (Stock No: 000664). *Abcc1^-/-^* (KO) and *Abcc1^+/+^* (WT) mice were generated from heterozygous crosses. Mice were born at expected Mendelian ratios and were genotyped by Polymerase Chain Reaction (PCR) analysis of genomic ear clip DNA using specific primers flanking exon 3 and part of exon 2 (P1 ’GTTTGAGCCACTCTCTCTGG‘, P2 ’GTGTTAAGCCGATGAGCAATC‘ and P3 ’CCTTCTATCGCCTTCTTGACG‘) as described previously (Lorico, et al. 1997; Nixon et al. 2016).

All experiments were performed in adult (> 8-week-old) male mice. Mice were maintained in individually ventilated cages in groups (3-5) at 21 °C with a light-dark cycle of 12h (lights on from 0700h to 1900h). Food and water were available *ad-libitum.* Mice were administered either control (chow) diet (2.71% kcal from fat, RM1(E) 801002, Special Diet Services) or high-fat diet (HFD; 58% kcals from fat plus sucrose, D12331, Research Diets) for up to 9 weeks. Body weight was measured weekly at the same time of the day in each group (AM). Mice were culled between 0900h and 1130h by decapitation, and trunk blood was collected in EDTA-coated microcentrifuge tubes and subjected to centrifugation (10,000 g, 5 min) to obtain plasma. All the procedures were performed under UK Home Office license and approved by the University of Edinburgh, Bioresearch & Veterinary Services (BVS).

### Physiological measurements

Insulin and Glucose Tolerance Tests were performed in animals following a 6 h fast (0900h to 1500h) on week 7 and week 8 of diet respectively. For Insulin Tolerance Test (ITT), insulin (0.75 U/Kg; Cat num. I9278, Sigma -Aldrich) was administered by intraperitoneal (IP) injection and blood glucose measured by glucometer (Accu-Check) in samples from tail venesection at 15-, 30-, 60-, 90- and 120-min post-injection. For Glucose Tolerance Test (GTT), glucose (1 g/Kg; Cat num. 50-99-7, Sigma-Aldrich) was administered by IP injection and blood glucose measured by glucometer as above at 15-, 30-, 60-, 90- and 120-min post-injection. Fasting insulin levels were assessed on plasma samples obtained at ‘0-min’ time point of ITT, and quantified by ELISA (Merck, EZRMI-13K). Diurnal sampling of blood for glucocorticoid profiling was performed (0800h and 2000h) by tail venesection and collected in EDTA-coated capillary blood tubes (Microvette), prior to analysis of steroids in plasma by liquid chromatography tandem mass-spectrometry (LC-MS/MS). ACTH levels were evaluated on plasma samples (trunk blood) by ELISA assay (MD Bioproducts, M046006). Homeostatic model assessment of insulin resistance (HOMA-IR) was calculated by using the formula (Fasting insulin (mg/dL) × Fasting glucose (mmol/L)/22.5.

### Western Blotting

Tissue protein lysates were prepared in RIPA lysis extraction buffer, supplemented with Halt^TM^, phosphatase and protease inhibitors (Thermo Scientific^TM^). Samples were disrupted using a TissueLyser II (QIAGEN) and 5mm stainless-steel beads (30Hz, 3 cycles of 15 sec). Protein concentration was quantified using bicinchoninic acid (BCA) assay (Thermo Scientific^TM^). Extracted proteins (25 µg) were resolved by SDS-PAGE, using Criterion TGX Precast Protein Gels 4-20% (Bio-Rad) under reducing and denaturing conditions. Proteins were transferred to nitrocellulose membranes using Trans-Blot® Turbo™ Blotting System (Bio-Rad). Membranes were blocked with 5% skimmed milk (Scientific Laboratory Supplies) in Tris-buffered saline, and then subjected to Western blotting using antibodies against OXPHOS proteins (ab110413, dilution 1:1000), and GLUT4 (MA1-83191, dilution 1:1000). Primary antibodies were used at described dilutions in 3% bovine serum albumin (BSA; Sigma-Aldrich) in Tris-buffered saline with Tween 20 and incubated overnight (4°C). Secondary antibodies (IRDye 800CW or IRDye 680CW (LI-COR) (anti-mouse, rat, and rabbit IgGs) were used at 1:10,000 dilutions in 3% BSA solution in Tris-buffered saline and incubated for 1 hour at room temperature. Total protein staining was performed using Revert^TM^ (LI-COR, 926-11011). Detection of protein was performed using an Odyssey CLx Imaging system (LI-COR). Densitometric analyses were performed using Image Studio^TM^ Software (LI-COR).

### RNA-Isolation and Quantitative RT-PCR

Tissue disruption was carried out using a TissueLyser II (QIAGEN) and 5mm stainless-steel beads (30 Hz, 3 cycles of 15 sec). Total RNA was extracted from tissues using Aurum^TM^ Total RNA Fatty and Fibrous Tissue Kit (Bio-Rad). Contaminating DNA was removed by treating the samples with DNase I (Bio-Rad). 500 ng RNA was used for reverse transcription using iScript reagent (Bio-Rad), and the products were analysed by qRT-PCR (LightCycler480, Roche). The qRT-PCR was performed using SybrGreen (iTAQ, Bio-Rad). Primer sequences are provided in **Table 1**. All primers were previously calibrated and used at efficiencies between 90-110%. Data analysis was performed using the Pfaffl method (Pfaffl 2001).

**Table 1.**
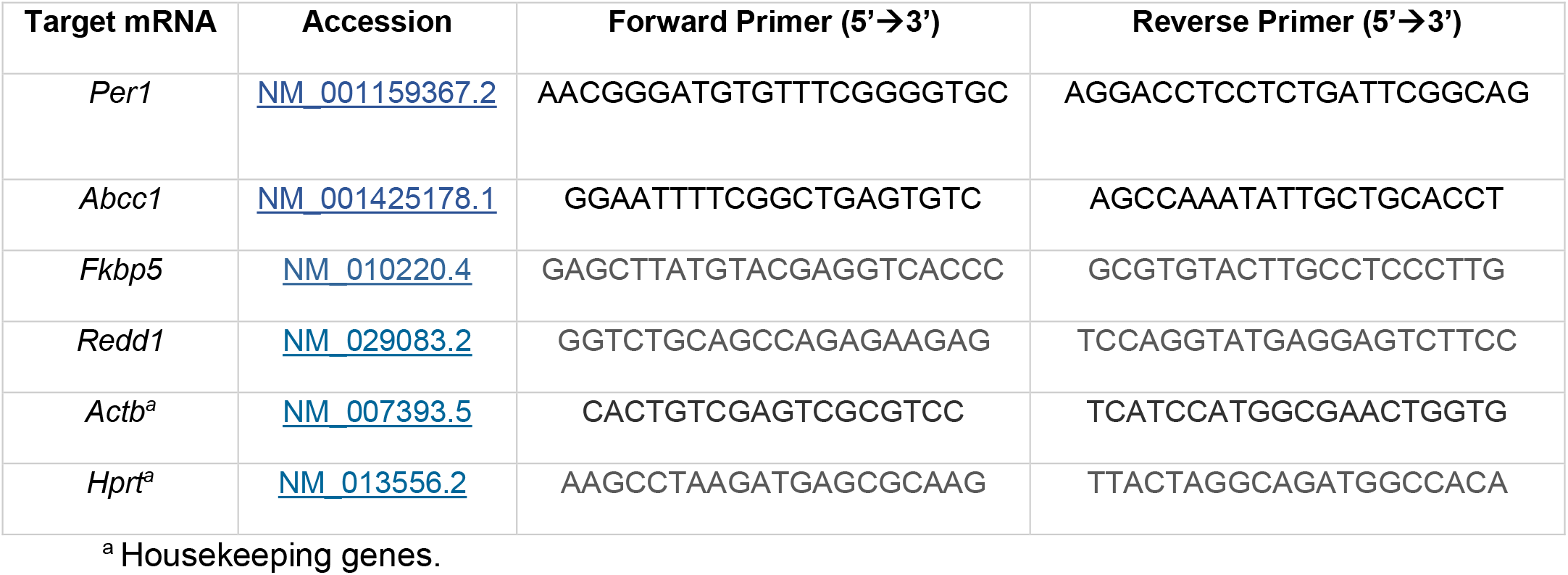
Forward and reverse primer sequenced for qRT-PCR.

### Steroid Profiling by liquid chromatography-tandem mass spectrometry (LC-MS/MS)

Glucocorticoids (corticosterone and 11-dehydrocorticosterone) were quantified in plasma derived from trunk blood and tail venesection, and in tissues as previously described (Bell et al. 2021). Briefly, steroid extraction was performed in samples of plasma (10 uL diurnal sampling and 100 µL trunk blood samples), liver (80-100 mg), gastrocnemius (40-50 mg) and sWAT (50-80 mg). Tissues were homogenised (Bead ruptor elite Bead Mill Homogenizer, Omni international) in acetonitrile with formic acid 0.1 %. A calibration standard curve (0.0025-500 ng/mL) was prepared and run alongside the samples. Homogenates were centrifugated and filtered through a Biotage Filter+ 96-well plate (0.22 µm), after which all samples and standards were enriched with d8-corticosterone as internal standard. The homogenate was then extracted using a ISOLUTE® PLD+ 96-well plate, for tissue samples (Biotage, Uppsala, Sweden) and a Microsolute® SLE 200 plate, for plasma (Biotage, Uppsala, Sweden), and eluted under positive pressure. Extracts were dried down under nitrogen at 40 °C and re-suspended for analysis by LC-MS/MS using a Waters I-Class UPLC connected to a QTrap 6500+ Mass Spectrometer (AB Sciex). Standards and samples were injected (20 μL) onto a Kinetex C18 column (150 × 2.1 mm and 2.6 μm; Phenomenex, #TN-1063) fitted with a 0.5 μm Ultra KrudKatcher (Phenomenex, #00F-4783-AN) at a flow rate of 0.3 mL/minute. The mobile phase system comprised water with 0.05 mM ammonium fluoride and methanol with 0.05 mM ammonium fluoride. The mass spectrometer was operated in positive ion electrospray ionisation mode using multiple reaction monitoring of steroids and internal standards. The instrumentation was operated using Analyst 1.6.3 (AB Sciex) and quantitative analysis of the data was carried out by least squares regression of the peak area ratio of the steroid to the corresponding internal standard with equal or 1/x weighting using MultiQuant software v3.0.3 (AB Sciex).

### Bulk RNA sequencing analysis

Samples (n=4 per experimental group) of gastrocnemius muscle and adipose tissue (sWAT) were processed for RNA isolation (as above). Total RNA was quantified using a Nanodrop spectrophotometer (Thermo Scientific^TM^) and integrity was assessed using an Agilent 2100 Bioanalyzer (Agilent Technologies Inc.) and Agilent RNA 6000 Nano kit. Library preparation and transcriptome sequencing was conducted by Novogene Co. Ltd.. cDNA libraries were sequenced using Illumina NovaSeq platform (Ilumina Inc). Bioinformatic analysis was performed by Fios Genomics (Edinburgh, UK). Quality of the data was assessed by the FastQC control tool. Reads were aligned to a mouse reference genome build GRCm39 using STAR aligner followed by calculation of alignment and mapping statistics. At least 87% of read pairs were uniquely mapped to one region of the genome.

### Proteomics

<u>Protein extraction:</u> Samples (n=4 per experimental group) of gastrocnemius (50-120 mg) muscle and subcutaneous white adipose tissue (70-230 mg) were homogenised in an extraction buffer (5% Sodium Dodecyl Sulfate (SDS) in 50mM Triethylammonium bicarbonate buffer (TEAB), pH 8.5) at sample to buffer ratio of 1:10, (w/v) using a Precellys homogenizer (5000-2x10) with bead in a ceramic vial (Precellys Lysing Kit, Tissue homogenizing CK mix). Following homogenization samples were centrifuged for 10min at 16,000g and supernatant was transferred into a clean low protein-binding vial then sonicated for 10 cycles with 30 sec on and 30 sec off per cycle. (Pico Sonicator Diagenode bioruptor). After sonication samples were centrifuged (16,000xg for 10min), the), supernatant was collected and a BCA assay performed.

S-Trap proteolytic digestion and liquid chromatography-mass spectrometry (LC-MS): The proteins were reduced with dithiothreitol and alkylated with iodoacetamide prior to tryptic digestion on S-TRAP (Protifi, USA) cartridges following manufacturer’s protocol. The resulting peptides were cleaned up using C18 stagetips. Purified peptides were separated over a 90 minutes gradient on an Aurora-25 cm column (IonOpticks Australia) using an UltiMate RSLCnano LC System (Dionex) coupled to a timsTOF FleX mass spectrometer via a CaptiveSpray ionization source. The gradient was delivered at a flow rate of 200 nL/min and washout was performed at 500 nL/min. The column temperature was set at 50°C. For DDA-PASEF acquisition, full scans were recorded from 100 to 1700 *m/z* spanning from 1.45 to 0.65 Vs/cm^2^ in the mobility (1/K0) dimension. Up to 10 PASEF MS/MS frames were performed on ion-mobility separated precursors, excluding singly charged ions which are fully segregated in the mobility dimension, with a threshold and target intensity of 1750 and 14,500 counts, respectively. Raw mass spectral data was processed using PEAKS Studio X-Pro Software (Bioinformatics LTD). Searches were performed against Uniprot mouse sequence database with MS1 precursor tolerance of 20 ppm and MS2 tolerance of 0.06 Da. Fully-tryptic digestion allowing one missed cleavage, fixed modification of cystine [+57.02] and oxidation of methionine and deamination of asparagine and glutamine were also specified for database search. Label-Free Quantitative analysis (LFQ) was performed with default parameters and with optional identification transfer enabled.

### Statistical Analysis

Results are expressed as mean ± SEM. Sample size (n=12 per group) was calculated based on the variability of a GTTs previously performed in C57Bl/6J mice under Chow and HFD conditions to detect differences of 20% (α=0.05 and Power=90%) in blood glucose. Downstream analyses (biomolecular) were performed in representative subset of the samples (n=3-8), detailed in each legend. Analyses were performed using Graph Pad Prism (Version 9.2.0, 2021 GraphPad Software LLC., San Diego, CA, USA). Comparisons between WT and KO mice on different diets were by Two-Way ANOVA, followed by a *post-hoc* test (*Tukey*). Comparisons of measurements over time were performed by Two-Way ANOVA with repeated measures, followed by a *post-hoc* test (*Sidak*), Normal distribution of the samples was assessed by Shapiro-Wilk test (*p* > 0.05), and if the dataset did not have a normal distribution, a non-parametrical test was used (Mann-Whitney U test or Kruskal-Wallis). Differences with a *p*-value < 0.05 were considered statistically significant.

## RESULTS

### In lean mice, *Abcc1* deficiency does not impact weight gain, glucose tolerance, or insulin resistance

To evaluate the role of *Abcc1* on the metabolic phenotype, *Abcc1* KO male mice (8-12 weeks old) and wild type (WT) littermates were fed either chow diet or high-fat diet (HFD; 58% kcal fat w/ sucrose) for 9 weeks. Mice with *Abcc1* deficiency did not differ in body weight from WT mice while receiving chow diet. (Figure 1A). Moreover, fat mass (gonadal and subcutaneous WAT) was not significantly different in KO (Figure 1B-E) compared to WT mice. However, brown adipose tissue (BAT) mass was significantly lower in KO vs WT mice (Figure 1F). Both insulin tolerance (Figure 1G-H, and Figure S1 A-B) and glucose tolerance (Figure 1I-J) were not different between KO and WT mice on chow diet.

**Figure 1.**
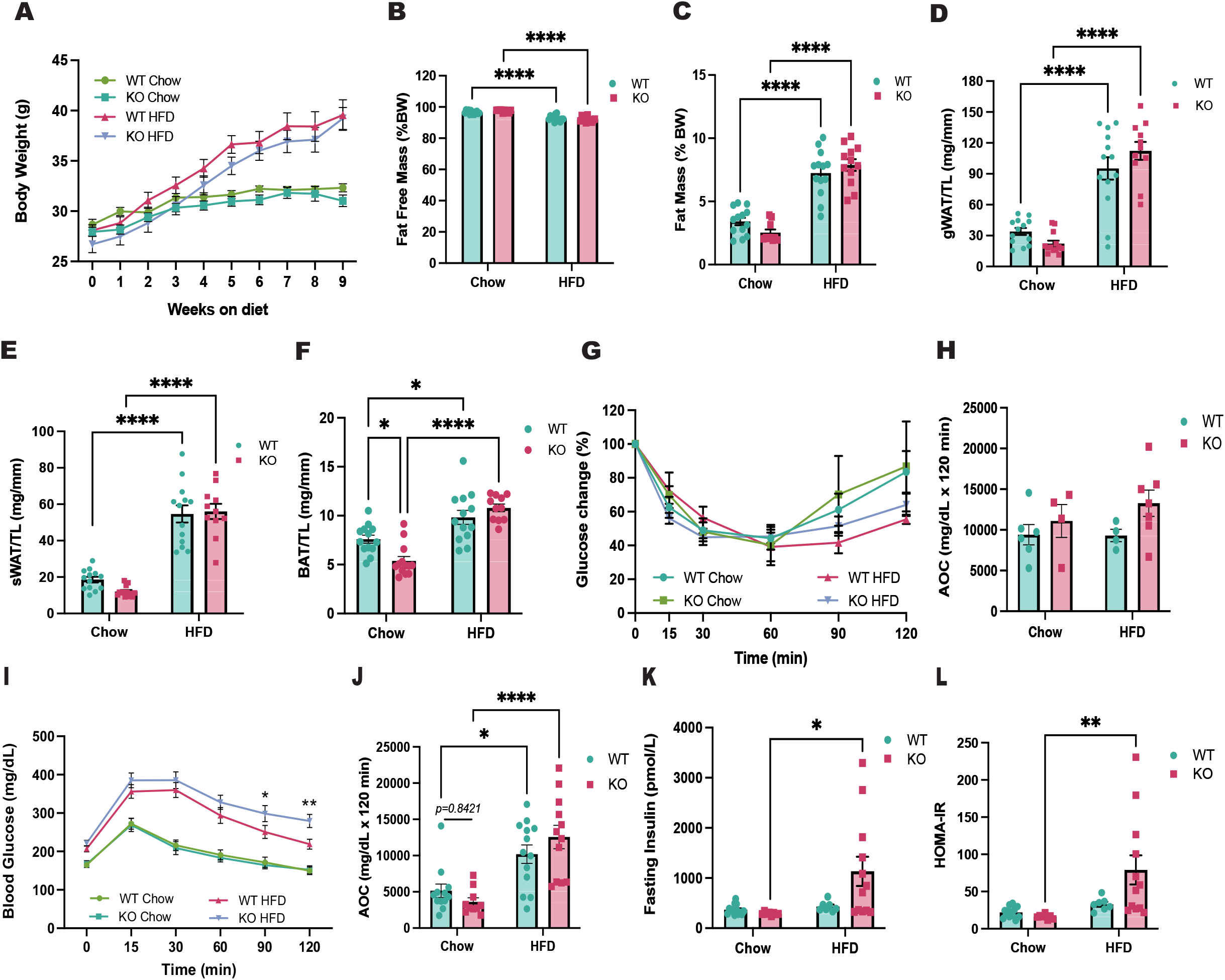
Comparison of metabolic profile between age matched *Abcc1* global knockout (KO) and wild-type littermates (WT) mice shows amplified insulin resistance in Abcc1 KO on high fat diet (HFD). Animals (12-13 per group) were fed either chow diet or HFD feeding started at 8- to 10-weeks of age and continued for 9 weeks. **(A)** Body weight, measured weekly, in WT and KO male mice while receiving chow and HFD. Data were analysed by a mixed effect model with Holm-Sidak’s multiple comparisons test. **(B)** Percentage (%) of fat free mass and fat mass **(C)**, with respect to total body weight, at the end of the study. **(D)** Weight of gonadal white adipose tissue (gWAT), **(E)** subcutaneous white adipose tissue (sWAT), and **(F)** brown adipose tissue (BAT), normalised by the length of the tibia (TL). Metabolic tests were performed after 7-8 weeks of chow or HFD, after 5 h fast. **(G)** Intraperitoneal insulin tolerance test (IP-ITT) performed after 7 weeks of dietary intervention in WT and KO mice, results shown as glucose change (percentage) from baseline, and **(H)** quantification of area over the curve (AOC) (n=4-7 animals per group). **(I)** Intraperitoneal glucose tolerance test (IP-GTT) performed after 8 weeks of dietary intervention in WT and KO mice, and **(J)** quantification of area over the curve (AOC) (n=8-13, animals per group). **(K)** Fasting insulin levels in WT and KO mice after 7 weeks of dietary intervention. **(L)** Homeostatic model assessment for insulin resistance (HOMA-IR). *p < 0.05, **p < 0.01, ***p < 0.001, ****p < 0.0001 by repeated measures ANOVA (A) and two-way ANOVA with Tukey’s multiple comparisons test. Data are expressed as mean ± SEM.

### In diet-induced obesity, *Abcc1* deficiency exacerbates glucose intolerance and insulin resistance

To determine the influence of HFD on *Abcc1*, we assessed mRNA levels in WAT and skeletal muscle. Our results showed similar levels of *Abcc1* in WT mice in chow versus HFD conditions in both tissues (Figure S1 C-D).

Under HFD-fed conditions, there were no differences between genotypes in measures of fat mass (Figure 1C) or adipose depot tissue weights, including BAT (Figure 1D-F). Insulin tolerance remained similar in WT and KO mice (Figure 1 G-H), but glucose intolerance with HFD was exacerbated in KO mice (Figure 1 I-J). Moreover, fasting insulin levels (Figure 1K) and HOMA-IR (Figure 1L) were significantly elevated in KO vs WT mice under HFD conditions.

### *Abcc1* deficiency increases tissue corticosterone in lean but not obese mice, and not independently of circulating levels

To determine if the metabolic phenotype in obese *Abcc1* deficient mice was a result of increased tissue glucocorticoid action in key metabolic tissues, we assessed systemic and tissue glucocorticoid concentrations under both chow and HFD conditions. Assessment of diurnal corticosterone from tail-vein plasma showed no differences between genotypes or diet (week 7, Figure 2A-B). On chow diet, tissue corticosterone levels were increased in adipose tissue (Figure 2C) and skeletal muscle (Figure 2D) of KO mice compared to WT controls. Unexpectedly, quantification of plasma corticosterone in trunk blood collected at the time of tissue collection (week 9, Figure 2E) revealed a striking similarity with tissue levels, with increased circulating corticosterone evident in KO mice compared to WT controls. These differences in tissue and plasma corticosterone between genotypes were abolished under HFD conditions.

**Figure 2.**
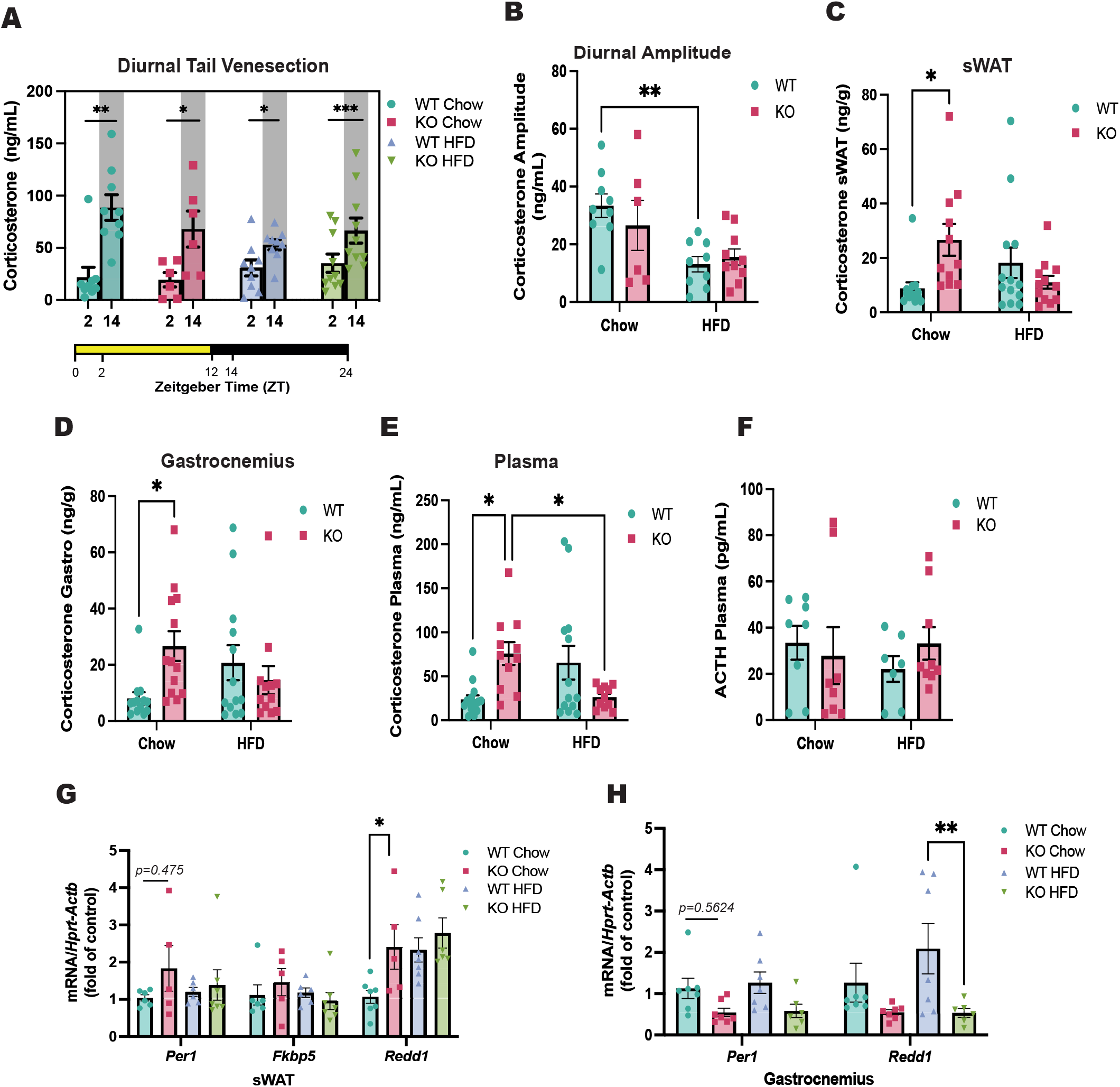
Deficiency of *Abcc1* induces accumulation of corticosterone in plasma, subcutaneous adipose tissue and gastrocnemius muscle in lean but not obese mice. Steroids levels were evaluated by LC-MS/MS in plasma and tissue samples of WT and KO male mice at week 7 and at the end of the study (week 9). **(A)** Corticosterone levels in plasma of mice obtained by tail venesection. Samples were collected at week 7 of the study, at 2 and 14 hours after light onset (Zeitgeber time). **(B)** Quantification of corticosterone diurnal amplitude in (A) (n=6-10, animals per group). Levels of corticosterone in terminal samples of **(C)** subcutaneous white adipose tissue, **(D)** gastrocnemius muscle, and **(E)** plasma (n=8-13, animals per group). **(F)** Plasma levels of adrenocorticotropic hormone (ACTH) (n=7-8, animals per group). **(G)** Evaluation of glucocorticoid-responsive genes (*Per1, Fkbp5 and Redd1*) in subcutaneous white adipose tissue, and **(H)** gastrocnemius muscle by qRT-PCR (n=6-7, animals per group). *p < 0.05, **p < 0.01. Data were analysed by Two-Way ANOVA with Tukey’s multiple comparisons test, and expressed as mean ± SEM.

To explore the basis for altered corticosterone levels, we measured plasma ACTH and infused adrenalectomised mice with corticosterone to measure steady state plasma concentrations and infer clearance of corticosterone. Neither plasma ACTH (Figure 2E) nor steady-state exogenous corticosterone levels (Figure S2 A-C) were different between genotypes under either dietary condition.

To determine if tissue glucocorticoid action mirrored the changes in corticosterone levels, we assessed several known glucocorticoid-responsive transcripts in adipose tissue (Figure 2G) and skeletal muscle (Figure 2H). In sWAT, *Abcc1* deficiency did not alter *Per1*, *Fkbp5 or Redd1* under HFD conditions, however *Redd1* was increased in KO animals under chow conditions. (Figure 2G). In skeletal muscle, neither *Per1* nor *Redd1* were elevated in KO mice and indeed *Redd1* was paradoxically lower in KO vs WT mice on HFD (Figure 2H).

In the absence of elevated tissue glucocorticoid levels and glucocorticoid-regulated transcripts in adipose or skeletal muscle in *Abcc1* deficient mice on HFD, we explored other mechanisms which might explain their adverse metabolic phenotype.

### Transcriptomic and proteomic analyses reveal differential responses to high-fat diet in adipose tissue from *Abcc1*-deficient mice

We performed bulk RNA sequencing (RNA-Seq) analysis in sWAT from WT and KO mice. Under both chow (Figure 3A) and HFD (Figure 3B) conditions, differentially expressed genes (DEGs) were not observed between genotypes following adjustment for multiple comparisons. However, when comparing the effect of HFD in each genotype (Figure 3C), KO mice displayed a strikingly heightened response to dietary intervention, with > 6000 unique DEGs compared to only 16 in WT mice.

**Figure 3.**
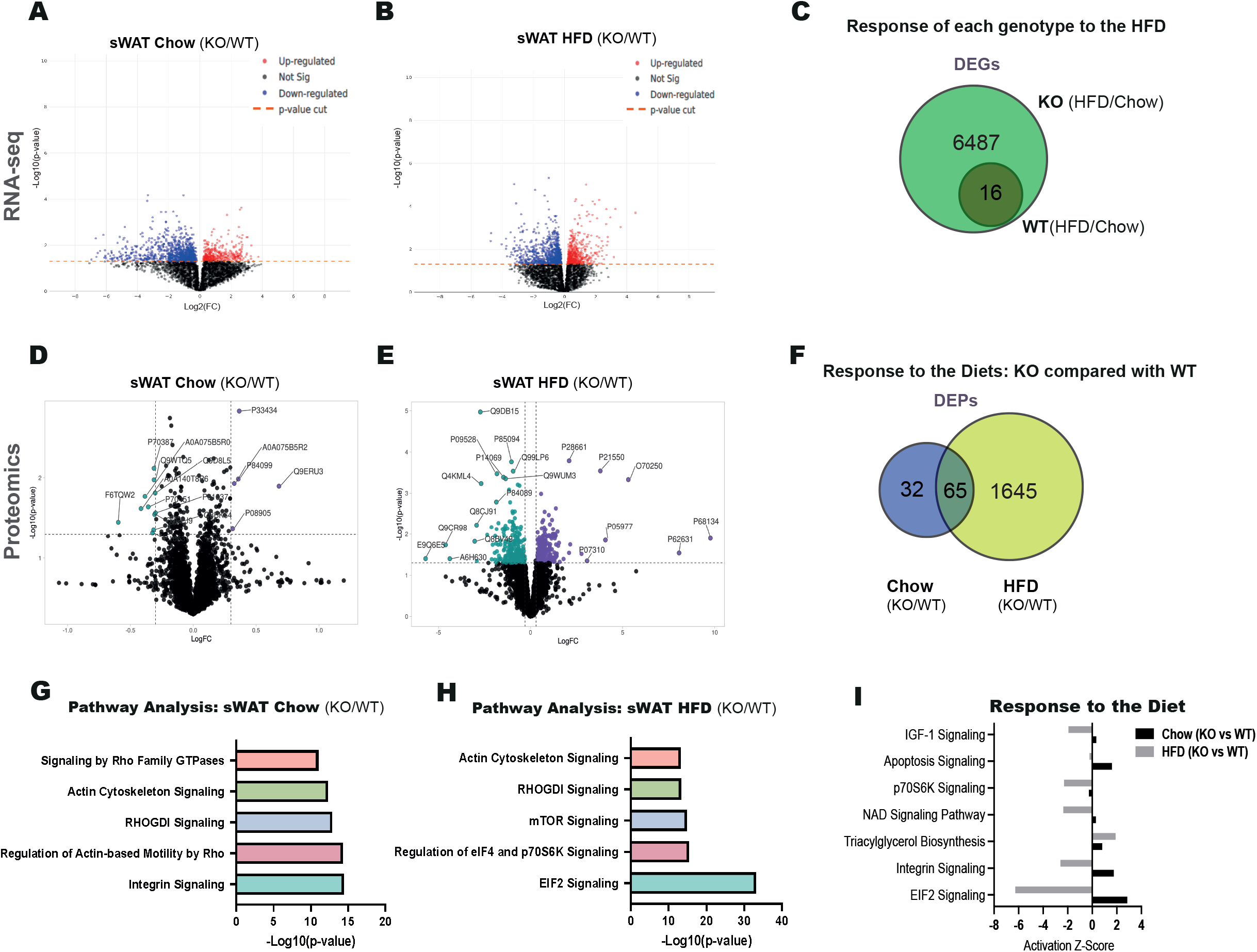
Transcriptomic and Proteomic analyses of subcutaneous adipose tissue reveal an amplified impact of HFD in *Abcc1* KO mice. **(A)** Volcano plot showing the differential expression analysis of all the genes identified in sWAT between wild-type (WT) and *Abcc1* KO mice fed with chow diet or **(B)** HFD during 9 weeks. **(C)** Venn diagram showing differential expressed genes (DEGs) of each genotype in response to the HFD (FC > 1.2 and adjusted *p-value* < 0.05). **(D)** Volcano plot showing the differential expression analysis of proteins in sWAT between wild-type and *Abcc1* knockout mice fed with control diet (chow) or **(E)** HFD for 9 weeks. Proteins differentially expressed are shown in purple (up-regulated) and green (down-regulated). **(F)** Venn diagram showing proteins with differential expression in KO compared with WT, in response to the diets**. (G)** Ingenuity pathway analysis (IPA) of differentially expressed proteins (DEPs) in sWAT of WT and *Abcc1* KO mice fed chow diet and **(H)** HFD, X-axis indicates – Log10 of the *p-value* and Y-axis indicates the corresponding canonical pathways. **(I)** Comparative analysis of differential activation on pathways identified in KO vs WT mice under chow and HFD conditions. X-axis indicates Z-Score explaining activation of the pathways in the Y-axis. Both omics analyses were performed in 4 animals per experimental group.

We also undertook a complementary proteomics approach. In contrast to the transcriptomic data, under both chow (Figure 3D) and HFD (Figure 3E) conditions, differentially expressed protein (DEPs) were identified when comparing between genotypes. Comparing KO vs WT mice, there were more genotype-dependent differences under HFD than under chow diet (1645 vs. 32 unique DEPs), (Figure 3F).

We also evaluated whether genes that are differentially expressed in KO mice are likely to reflect altered glucocorticoid signalling. We identified genes in WAT that have previously been shown to be regulated by glucocorticoids both in mice (following dexamethasone treatment) and in humans (inferred from trans-QTL analyses for genes associated with variation in plasma cortisol) (Bankier, et al. 2023). These included *Pkp2, Osmr, Phyh, Zc3h7b,* and *Me2*; none of these were differentially expressed when comparing sWAT gene expression in KO vs WT mice.

Ingenuity Pathway Analysis (IPA) was used to identify enriched biological networks and determine potential regulatory mechanisms. Here, the canonical pathways were obtained by overlapping the DEPs with the number of proteins covering canonical pathways, previously defined by the IPA database. The results show that in lean, chow-fed conditions, the top dysregulated pathways were related with tissue remodelling and traffic of vesicles (Figure 3G). In obese, HFD-fed mice, the most dysregulated pathways were associated to reticular stress, mTOR signalling and tissue remodelling (Figure 3H). A comparative analysis of the effect of the different diets was done in the KO mice (controlling for the WT expression). The IPA software performs the calculation of a z-score that gives a weight to the up/downregulation of proteins in sWAT, and can thus predict activation status of different pathways (Figure 3I). The results identified differential activation of signalling pathways related to EIF2, integrin, IGF-1, S6K, apoptosis, and NAD in KO mice under chow and HFD conditions.

### Transcriptomic and proteomic analyses identify impaired oxidative phosphorylation in skeletal muscle of *Abcc1*-deficient mice

Similar to sWAT, transcriptomic analysis of skeletal muscle did not reveal DEGs between *Abcc1* deficient and WT mice on either chow (Figure 4A) or HFD (Figure 4B). In contrast to the striking effect of HFD observed in the adipose tissue of *Abcc1* deficient mice (compared with WT), skeletal muscle showed a more modest response to HFD (Figure 4C). However, the transcriptional response to HFD was still exaggerated in KO mice compared with WT in skeletal muscle (5 DEGs in KO mice vs 0 in WT mice). As was observed in adipose tissue, proteomic analyses identified DEPs between genotypes under chow (Figure 4D) and HFD (Figure 4E) conditions, with a similar number of proteins differentially altered between genotypes in chow and HFD (357 vs. 470 respectively) (Figure 4F). IPA analysis revealed increased overlapping of proteins in pathways associated with reticular stress, mitochondrial dysfunction and oxidative phosphorylation in all the studies (Figure 4G-H). Comparative analysis using IPA showed predicted differential activation of pathways in KO mice under dietary conditions, with pathways related with TCA cycle, β-oxidation, and oxidative phosphorylation activated under chow diet, but inactivated under HFD conditions (Figure 4I).

**Figure 4.**
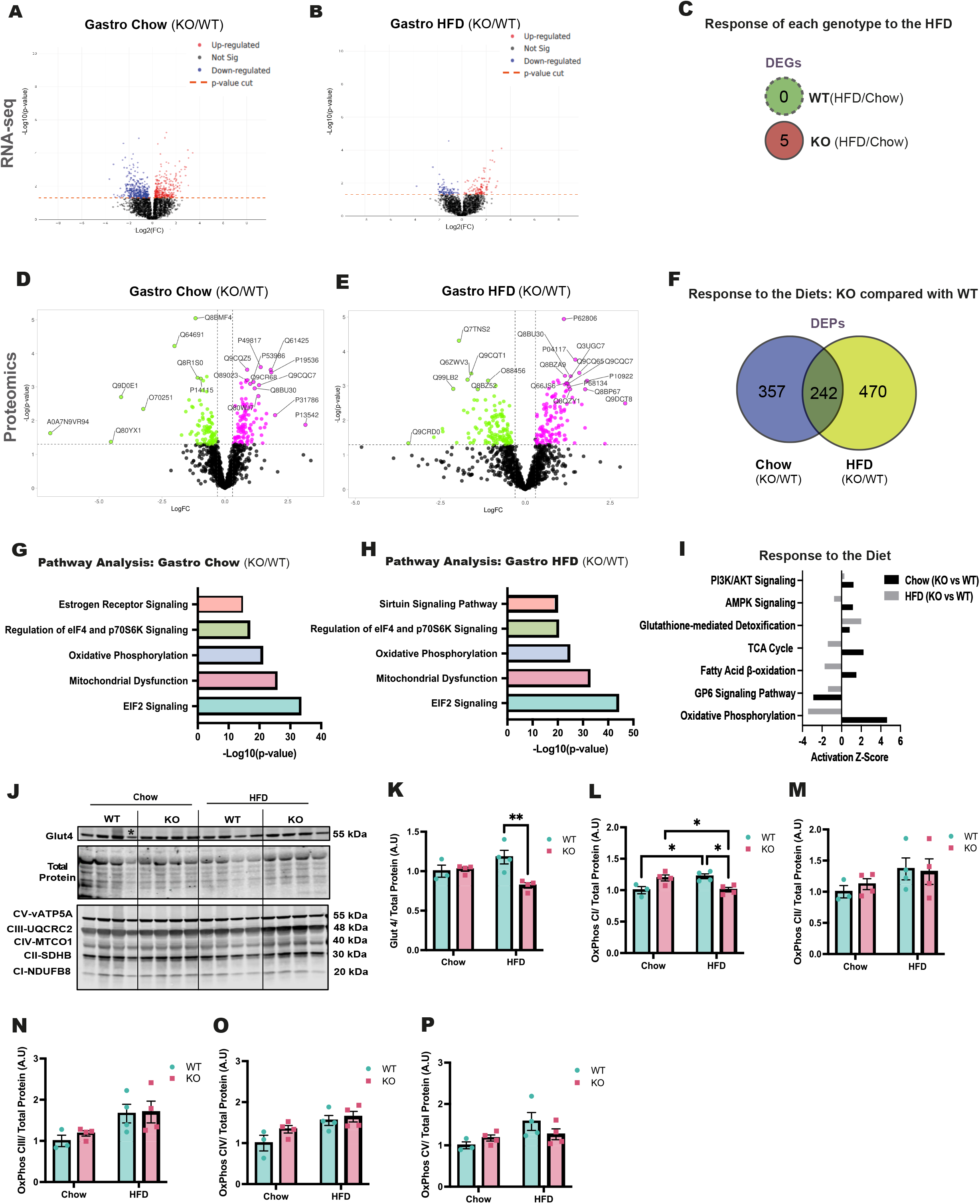
Proteomic and pathway analyses of *Abcc1* KO gastrocnemius muscle reveals impairment in oxidative phosphorylation in mice exposed to HFD. **A)** Volcano plot showing the differential expression analysis of all the genes identified in gastrocnemius muscle between WT and *Abcc1* KO mice fed with chow diet or **(B)** HFD for 9 weeks. **(C)** Venn diagram showing differential expressed genes (DEGs) of each genotype in response to the HFD (FC > 1.2 and adjusted *p-value* < 0.05). **(D)** Volcano plot showing the differential expression analysis in gastrocnemius between WT and *Abcc1* KO mice fed chow and **(E)** HFD during 9 weeks. Proteins differentially expressed are shown in fuchsia (up-regulated) and green (down-regulated). **(F)** Venn diagram showing proteins with differential expression in KO compared with WT, in response to the diets**. (G)** Ingenuity pathway analysis (IPA) of differentially expressed proteins in gastrocnemius muscle of wild-type and Abcc1 knockout mice fed with control (chow) diet or **(H)** HFD, X-axis indicates – Log10 of the p-value and Y-axis indicates the corresponding canonical pathways. **(I)** Comparative analysis of differential activation on pathways identified in KO vs WT mice under chow and HFD conditions. X-axis indicates Z-Score explaining activation of the pathways in the Y-axis. The omics analyses were performed in 4 animals per experimental group. **(J)** Western blot analysis (gastrocnemius) and (H-M) densitometric quantification of **(K)** Glut4 and OXPHOS: **(L)** Complex I, **(M)** Complex II, **(N)** Complex III, **(O)** Complex IV, **(P)** Complex V. Data was normalised by the staining of total proteins (n=3-4, animals per group). (*) Rout Method (Q=1%) was used to identified outliers. *p < 0.05, **p < 0.01. Data were evaluated by Two-Way ANOVA with Tukey’s multiple comparisons test, and are expressed as mean ± SEM.

The proteomic analysis showed changes in glycolysis and oxidative phosphorylation (Figure 4I). To understand if the dysregulation of these processes could have an impact on the metabolic phenotype of Abcc1 deficient mice, we went on to validate our results. We assessed the protein levels of components of oxidative phosphorylation complex machinery (OXPHOS), and Glut4, a key glucose transporter in skeletal muscle, to assess a potential dysregulation at the beginning of the glucose oxidation (Figure 4J-P). Western blots revealed decreased levels of Glut4 and Complex I in KO mice under HFD.

## DISCUSSION

Our prior work shows that *Abcc1* acts as a transporter of exogenously administered glucocorticoid in adipose tissue (Nixon et al. 2016). Here, we tested whether *Abcc1* has a role as a transporter of endogenous glucocorticoid in adipose tissue and skeletal muscle, and whether it influences adiposity and glucose metabolism. Although tissue glucocorticoid levels were elevated with *Abcc1* deficiency, this could not be dissociated from simultaneous differences in plasma glucocorticoid levels and was not sustained when mice were given a high fat diet. Moreover, analysis of transcriptomic and proteomic data in adipose tissue and skeletal muscle did not provide evidence for enrichment of glucocorticoid-responsive genes amongst those which differed with *Abcc1* deficiency. However, despite the lack of evidence for enhanced glucocorticoid action, *Abcc1* deficiency caused metabolic dysfunction and an exaggerated transcriptional and proteomic response to HFD. This suggests that, in contrast with our hypothesis, there is a protective metabolic effect of *Abcc1* which is glucocorticoid-independent.

We observed changes in plasma corticosterone with *Abcc1* deficiency in terminal trunk blood samples in lean mice but not in diurnal tail-nick samples. This suggests a greater response to stress during sacrifice in *Abcc1*-deficient mice. Previous studies suggested that ABCC1 influences glucocorticoid clearance (Livingstone, et al. 2017; Livingstone, et al. 2014), but we did not find evidence for this when we infused corticosterone in adrenalectomised mice. There is also evidence that ABCC1 influences HPA axis negative feedback in humans (Kyle et al. 2022) but we did not find differences in plasma ACTH to substantiate a central activation of the HPA axis, albeit that measurements of ACTH are labile and hence insensitive to assess changes in the HPA axis (Donegan, et al. 2019; Toprak, et al. 2016; Wu and Xu 2017). This unexpected phenomenon of higher plasma corticosterone during terminal sampling represents a confounder in the interpretation of endogenous tissue corticosterone levels in lean *Abcc1*-deficient mice that we have been unable to overcome. However, this effect appears to be over-ridden by the well-documented effect of HFD to alter HPA axis function (Morton, et al. 2005). The absence of elevated tissue glucocorticoid levels in *Abcc1*-deficient mice on HFD, along with the evidence that HFD did not alter *Abcc1* expression in our experiments, suggests that ABCC1 does not have a potent effect on tissue levels of endogenous corticosterone.

Even in the absence of altered tissue glucocorticoid levels, or of changes in adiposity, metabolic assessments revealed impaired glucose tolerance and hyperinsulinemia in *Abcc1* deficient mice under HFD. In light of this paradox, we sought to study key differences in the transcriptome and proteome in the adipose tissue and skeletal muscle of *Abcc1* deficient mice, which may be independent of glucocorticoid action and might mediate the adverse metabolic response (Calejman, et al. 2022; Cottam, et al. 2022). It is worthy of note that bulk RNA-seq can have some pitfalls when the heterogeneity of the tissues is high (Denninger, et al. 2022; Li and Wang 2021; Noureen, et al. 2022). An expansion of adipose tissue due to HFD exposure has been shown not only to change the cell profile in adipose tissue by increasing proinflammatory cells, but also in skeletal muscle (Chait and den Hartigh 2020; Fuster, et al. 2016; Miranda, et al. 2023; Morgan, et al. 2020).

Despite these limitations, in white adipose tissue, both mRNA and protein profiling revealed that *Abcc1* deficiency amplifies the response to HFD. Key signalling pathways that were differentially regulated include extracellular matrix (ECM) and reticular stress. A link between ECM and ABCC1 was described on HT-29 cells, where the ECM from tumour cells was able to upregulate *ABCC1*, increasing their chemoresistance capacity (Hoshiba and Tanaka 2016). We also observed the Eukaryotic Initiation Factor 2 (eIF2) signalling pathway to be less active in *Abcc1*-deficient mice under HFD). eIF2 is a key protein regulating reticular stress, a process that has been demonstrated to be a major contributor to the metabolic dysfunction induced by obesity (Han and Kaufman 2016; Li, et al. 2020).

In skeletal muscle, the proteomic analysis shed light on the pathways related to glucose homeostasis that could explain the impairment of glucose tolerance and hyperinsulinemia on the *Abcc1*-deficient mice receiving HFD. These included mitochondrial dysfunction, oxidative phosphorylation and reticular stress, all fundamental to metabolism (Antonopoulos and Tousoulis 2017; Rani, et al. 2016; Zhu, et al. 2022). Supporting the inference of impaired oxidative phosphorylation in skeletal muscle with *Abcc1* deficiency, our assessment of the OXPHOS complex identifying decreased protein levels in Complex I, the initiator of the respiratory chain (Sharma, et al. 2009). Further, diminished levels of Glut4 in skeletal muscle of KO mice under HFD, could contribute to the impairment of glucose tolerance in these mice. Added to the downregulation of NADH dehydrogenase or Complex I, in *Abcc1* KO mice under HFD, suggests alterations in the initiation of the electron transport chain, potentially having an impact on the entire process of oxidative phosphorylation. Overall, these represent a plausible pathway that could mediate the observed metabolic effects of *Abcc1*.

A limitation of our studies is that we focused on subcutaneous WAT, in line with previous work demonstrating a role for ABCC1 in modulating exogenous glucocorticoid levels (Nixon et al. 2016). We did observe a reduction in brown adipose tissue (BAT) weight in lean ABCC1-deficient mice but not in mice fed a HFD. We have not pursued this observation further here, although it is notable that a recent study showed BAT as the most sensitive adipose tissue depot during acute (1 week) corticosterone administration (Harvey, et al. 2023).

We have not determined the mechanism by which ABCC1 would influence the metabolic pathways that are implicated in white adipose tissue and skeletal muscle. ABCC1 is a multidrug resistance protein that acts on the efflux not only of glucocorticoids but also a variety of xenobiotics (Cole 2014; Devine et al. 2023). Endogenous substrates of ABCC1 include estradiol-17β-glucuronate (Cole 2014; Devine et al. 2023; Slot, et al. 2008), proinflammatory molecules (LTC4), antioxidants (GSH) and signalling lipids (SP1) (Cole 2014; Devine et al. 2023; Nieuwenhuis, et al. 2009). Many of these substrates are altered in in obesity (Antonopoulos and Tousoulis 2017; Cottam et al. 2022; Fuster et al. 2016) and could mediate the worsened metabolic profile in *Abcc1-*deficient mice under HFD. Our findings of a novel role for ABCC1 in limiting the adverse metabolic effects of obesity should stimulate further investigation of the various substrates of ABCC1 and their transmembrane transport in obesity.

## DATA AVAILAILITY

Data available upon request or from the University of Edinburgh Data Store https://datashare.ed.ac.uk

## SUPPLEMENTARY MATERIALS

Figure S1 and S2 are attached in a PDF file.

## DECLARATION OF INTEREST

All the authors declare no conflict of interests, financial or otherwise.

## FUNDING

Supported by a Wellcome Senior Investigator Award (to B.R.W.) and an Early Career Grant from the Society for Endocrinology (to E.V.). T.M.W. would like to acknowledge support from the BBSRC ISP1 BBS/E/RL/230001C and core capability funding.

## AUTHOR CONTRIBUTION STATEMENT

E.V., B.R.W and M.N. conceived and designed the studies; E.V., A.M., R.A.M., L.I and J.A. performed experiments; E.V, R.A.K. and D.K. analysed data; E.V., B.R.W and M.N. interpreted results; E.V. prepared figures and drafted manuscript; E.V., B.R.W., R.A.M., N.M., R.A., R.S., T.M.W. and M.N. edited and revised manuscript; all authors approved the final version of manuscript.

## ACKNOWLEDGEMENTS

The authors thank the Mass Spectrometry Core, at the Queen’s Medical Research Institute (QMRI), University of Edinburgh for the skilled help and support. We thank Dr Sean Bankier for his guidance on the glucocorticoid-regulated gene network.

## Notes

### Competing Interest Statement

The authors have declared no competing interest.

