## Supplementary Materials for "ATP-Binding Cassette Family C member 1 constrains metabolic responses to high-fat diet in male mice"

### MATERIALS AND METHODS

#### Adrenalectomy and Mini-pump Insertion

Adult male mice (8-10- weeks old, n=7-8 per group), were bilaterally adrenalectomised under anaesthesia and the subcutaneous mini-pump inserted dorsally at the same procedure using the same opening. Muscle wall was closed with absorbable sutures and skin incisions closed with stainless steel clips. Mice were daily monitored and weight after surgery, to ensure recovery. Mice were single housed animals after surgery and 0.9% saline drinking water was provided.

Pump details: osmotic mini-pumps (Alzet #2001) containing steroid (corticosterone and cortisol; 10.4mg/ml each, in propylene glycol) was prepared the day before the study and primed in saline solution overnight at 37°C. The saline priming solution was changed an hour before pump insertion. The pump was in situ for 7 days. Mice were culled between 9:00 to 11:30AM by decapitation, and the trunk blood and tissues were collected to perform steroid measurements. All the procedures were performed under UK Home Office license and approved by the University of Edinburgh, Bioresearch & Veterinary Services (BVS).

### FIGURE LEGENDS

**Supplementary Figure 1. Insulin tolerance test raw glucose values, and *Abcc1* expression in tissues.**

Animals were weaned onto chow diet from 4-weeks of age and either chow diet or HFD feeding started at 8- to 10-weeks of age and continued for 9 weeks. **(A)** Intraperitoneal insulin tolerance test (IP-ITT) performed at week 7 of the study in WT and KO mice fed with chow diet and HFD. **(B)** quantification of area under the curve (AUC) in (A) (n=4-7 animals per group). **(C)** Evaluation of *Abcc1* gene expression in subcutaneous white adipose tissue, and **(D)** tibialis anterior muscle by qRT-PCR (n=5-6, animals per group). Statistical analysis was done by Two-Way ANOVA with repeated measures (A-B), followed by a *post-hoc* test (*Sidak*). A *p-value* > 0.05 was considered significant. Data are expressed as mean ± SEM.

**Supplementary Figure 2. Glucocorticoid Clearance**

Animals were weaned onto chow diet from 4-weeks of age and chow diet started at 8- to 10-weeks of age. Abcc1 KO mice and WT littermates (7-8 per group) were bilaterally adrenalectomized and the subcutaneous mini-pump inserted to deliver corticosterone and cortisol (10.4 mg/mL each) during 7 days. Steroids levels were evaluated in terminal samples by LC-MS/MS. in plasma (A) and tissue samples. **(A)** Levels of corticosterone in plasma (ng/mL), (**C)** subcutaneous white adipose tissue (ng/g), and **(C)** gastrocnemius muscle (ng/g). *$p$-value $< 0.05$, by Student's t-test between WT vs. KO mice. Data are expressed as mean ± SEM.
