## Supplementary Figures for "ATP-Binding Cassette Family C member 1 constrains metabolic responses to high-fat diet in male mice"

#### Figure S1. Insulin tolerance test and Abcc1 expression in tissues

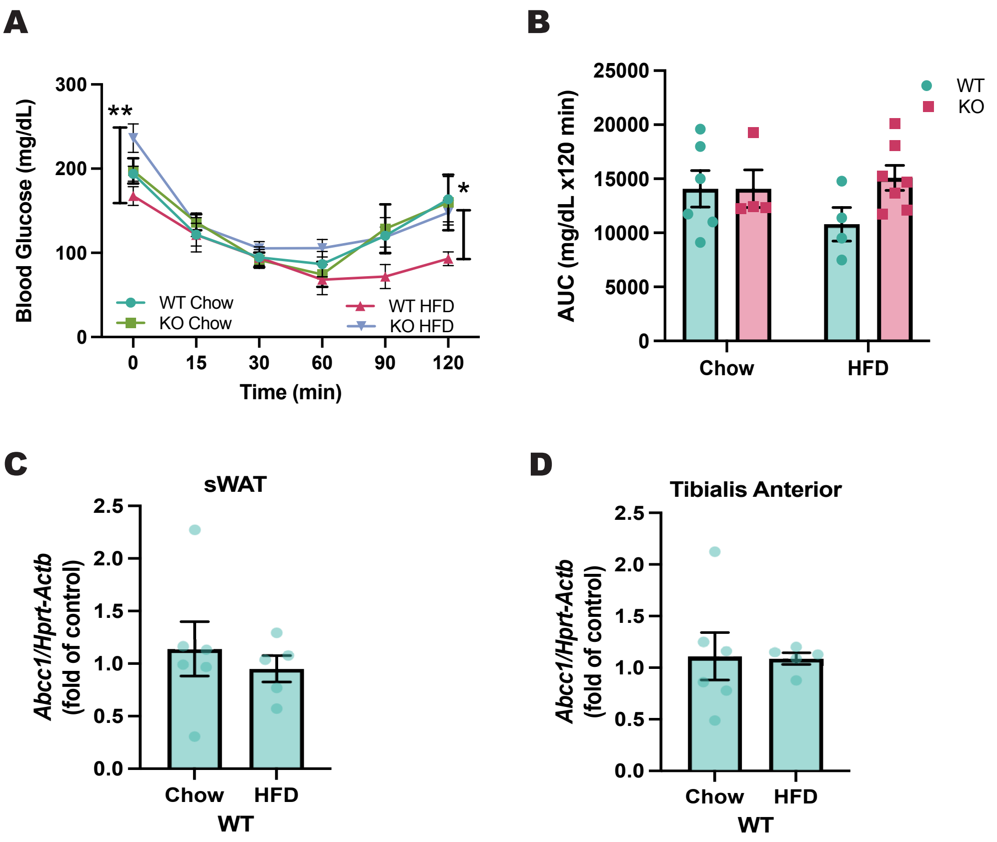

### Figure S2. Glucocorticoid Clearance

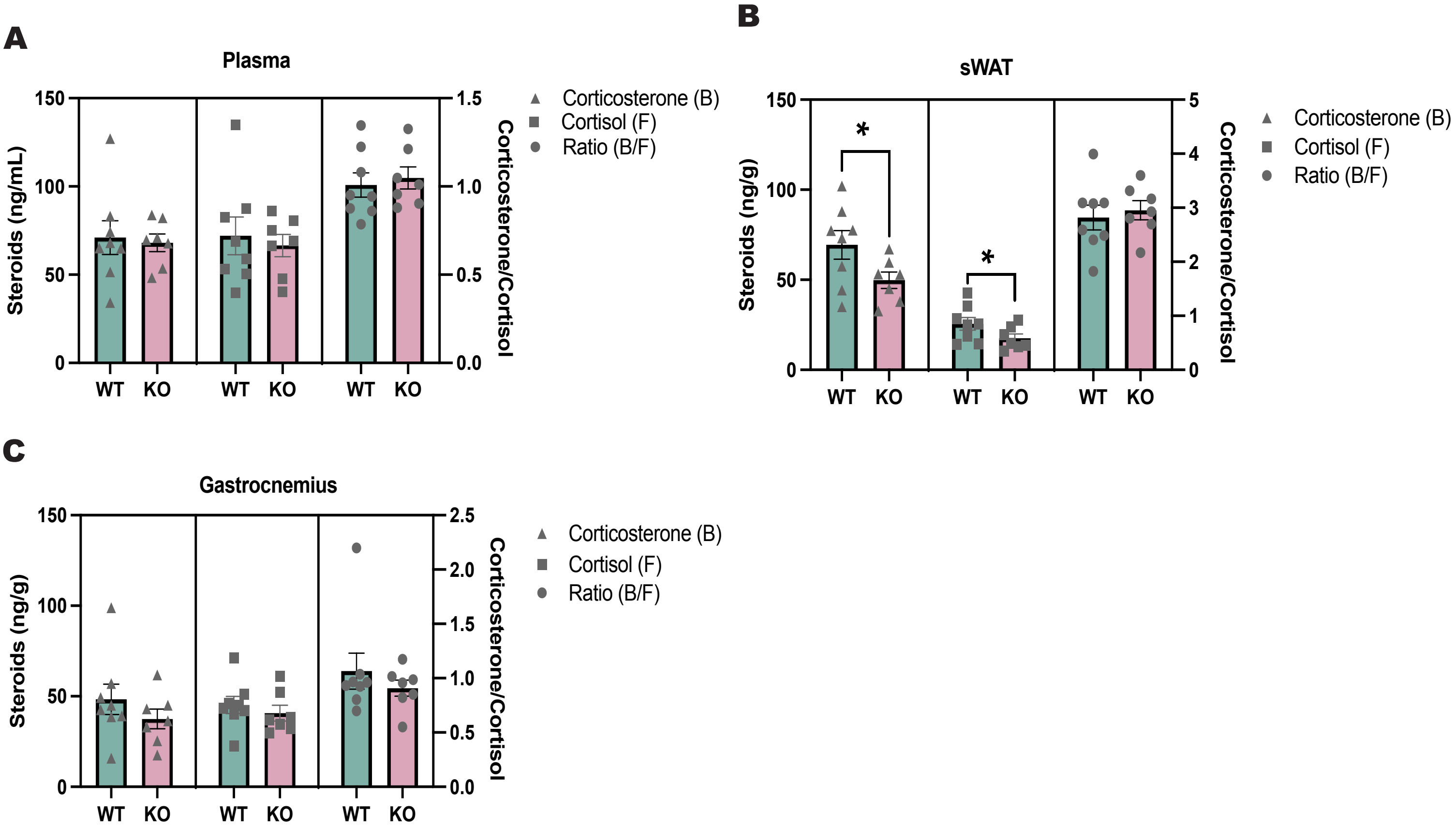
